# Distribution of iridescent colours in hummingbird communities results from the interplay between selection for camouflage and communication

**DOI:** 10.1101/586362

**Authors:** Hugo Gruson, Marianne Elias, Juan L. Parra, Christine Andraud, Serge Berthier, Claire Doutrelant, Doris Gomez

## Abstract

Identification errors between closely related, co-occurring, species may lead to misdirected social interactions such as costly interbreeding or misdirected aggression. This selects for divergence in traits involved in species identification among co-occurring species, resulting from character displacement. On the other hand, predation may select for crypsis, potentially leading co-occurring species that share the same environment and predators to have a similar appearance. However, few studies have explored how these antagonistic processes influence colour at the community level. Here, we assess colour clustering and overdispersion in 189 hummingbird communities, tallying 112 species, across Ecuador and suggest possible evolutionary mechanisms at stake by controlling for species phylogenetic relatedness. In hummingbirds, most colours are iridescent structural colours, defined as colours that change with the illumination or observation angle. Because small variations in the underlying structures can have dramatic effects on the resulting colours and because iridescent structures can produce virtually any hue and brightness, we expect iridescent colours to respond finely to selective pressures. Moreover, we predict that hue angular dependence – a specific aspect of iridescent colours – may be used as an additional channel for species recognition. In our hummingbird assemblages in Ecuador, we find support for colour overdispersion in ventral and facial patches at the community level even after controlling for the phylogeny, especially on iridescence-related traits, suggesting character displacement among co-occurring species. We also find colour clustering at the community level on dorsal patches, suspected to be involved in camouflage, suggesting that the same cryptic colours are selected among co-occurring species.

This article has been peer-reviewed and recommended by *Peer Community In Evolutionary Biology*

## Introduction

Colour is a complex communication channel widespread among various taxa and involved in many ecological and evolutionary processes [7]. It can be described by multiple variables, including hue (colour in its common sense, such as red, green, blue, etc.) and brightness (average level of grey of a colour, i.e. whether the object is light or dark). Colours can be produced by two non-mutually exclusive means: pigmentary colours are produced by the selective absorption of incoming light by pigments, while structural colours are produced by the interaction of incoming light with nanostructures, causing diffraction, interferences or scattering [68]. Among structural colours, iridescent colours are characterised by a shift in hue with changes in illumination or observation angle [88]. Iridescent colours are found in many bird families such as Anatidae (ducks) Phasianidae (fowls), Sturnidae (starlings), or Trochilidae (hummingbirds), and thought to be involved in numerous adaptations [24]. But evolution of iridescent colours at the community level remains poorly understood. Yet, evolutionary patterns of iridescent colours, which remain poorly studied and understood, may differ from that of non-iridescent colours. Indeed, as opposed to other types of colours, iridescent colours can produce virtually any hue and are expected to respond more readily and finely to selection, because large changes of hue can be achieved by small changes in the underlying structures [72]. They can also result in directional colours only seen at specific angles, as well as highly reflective colours [65].

Because colours are involved in many different ecological processes, they are subject to multiple selection pressures, often with opposite effects [33]. Colour may indeed increase or decrease detectability of an animal depending on the colour constrast with its surroundings. In particular, colour can reduce predation risk via crypsis or aposematism or serve as a means of species identification. In this case, two opposite evolutionary forces act on colours: (i) On the one hand, species living in the same environment are likely experiencing similar selective pressures, such as predation. The environment is characterised by ambient light and vegetation, which both influence greatly which colours are poorly detectable and which colours are highly detectable [29, 32]. We thus expect co-occurring species to harbour the same, poorly detectable, colours as this would decrease the risk of being detected by predators, thereby causing a clustering pattern in colouration at the community level, all else being equal. This colour clustering can result from convergence between sympatric species (evolutionary process), from environmental filtering (ecological process), i.e. species sorting locally according to the traits they harbour, or a mixture of the two (detailed in table 1). (ii) On the other hand, sympatric closely-related species are more likely to face problems of species recognition, eventually resulting in reproductive interference - a phenomenon where an individual courts or mates with individuals of another species, producing no offspring or low fertility hybrids, leading to costly interbreeding [38]. Species misidentification can also lead to misdirected aggression and costly fighting when individuals compete over resources or territories. Hence, any feature that would enhance species recognition is expected to be selected for. In this context, closely related species living in sympatry should be under strong selective pressure to diverge in traits involved in communication, if divergence enhances species recognition. Divergence can result from a process called character displacement (RCD for reproductive character displacement, ACD for agonistic character displacement; evolutionary process) [8, 9, 37] or from species sorting (ecological process). For ACD, it is worth noting that traits are expected to diverge only in case of moderate ecological competition, whereas they should converge in case of high competition [37, 86]. Multiple empirical studies have shown character displacement for songs (e.g. Gerhardt [31] in frogs and Grant and Grant [35] in birds), or olfactory signals [3]. However, fewer studies have looked at divergence in colour patterns (but see Doutrelant, Paquet, Renoult, Grégoire, Crochet, and Covas [25], Hemingson, Cowman, Hodge, and Bellwood [44], Lukhtanov, Kandul, Plotkin, Dantchenko, Haig, and Pierce [50], Martin, Montgomerie, and Lougheed [53], Naisbit, Jiggins, and Mallet [61], and Sætre, Moum, Bureš, Král, Adamjan, and Moreno [74]). Almost all these studies were at the species level, and at best involved comparison between closely related species. Many of them also did not use objective spectrometry measurements and instead relied on human vision, which did not allow them to analyse colours as perceived by the intended receiver, in the case of this study: birds [6, 16, 27, 59].

**Table 1.**
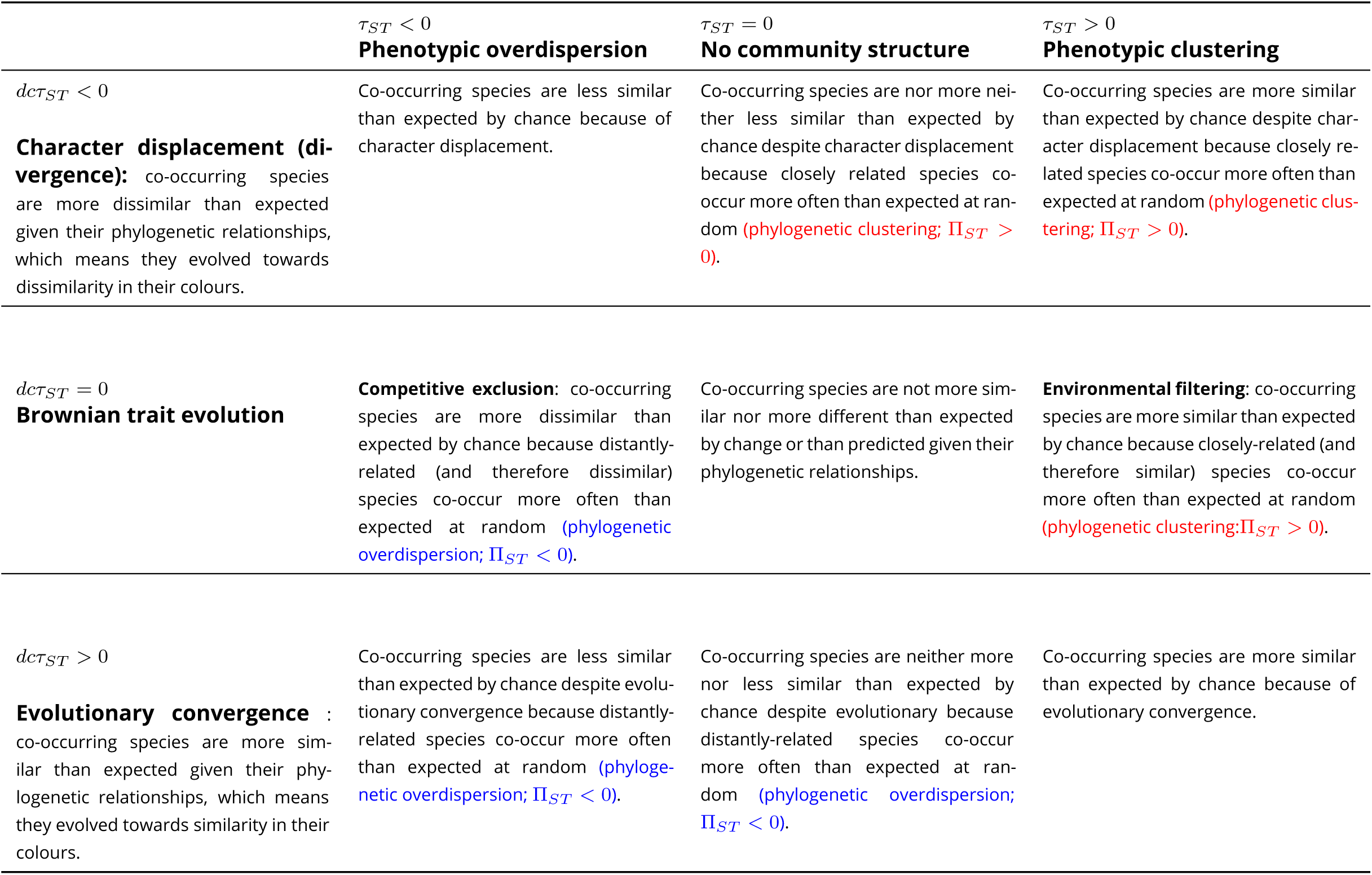
Summary of the different evolutionary and ecological scenarios and their results in terms of values of *τ*_*ST*_ and decoupled *dcτ*_*ST*_.

In birds, it has been shown that colouration is under different selective pressures depending on the body patch location: dorsal patches, which are exposed to aerial predators, are mainly involved in camouflage while ventral and facial patches are mainly involved in communication [21, 33]. In this study, we test this hypothesis for iridescent colours at the community level by looking at phenotypic structure in hummingbird local assemblages across different body parts. Hummingbirds are an interesting study system to test this hypothesis as various published accounts of sexual displays and aggressive encounters among hummingbirds have made clear that certain feather patches such as the crown and throat are consistently used during these displays [46, 75–78]. On the other hand, colours displayed on the dorsal side of hummingbirds tend to resemble background colours and thus have been suggested to be cryptic [70]. Accordingly, we predict that co-occurring hummingbird species should display similar hues on dorsal patches, leading to phenotypic clustering of hues (i.e. co-occurring species are more similar than expected by chance, prediction 1) and different hues on ventral patches, resulting in a phenotypic overdispersion pattern (i.e. co-occurring species are more dissimilar than expected by chance, prediction 2). For brightness, we can formulate two alternative predictions: on the one hand, it might evolve in the same way as hue, also because of reproductive character displacement and selection for camouflage, leading to the same outcome as for hue (prediction 3, equivalent to predictions 1 and 2 but for brightness). On the other hand, because brightness level positively correlates with signal conspicuousness, poorly detectable signals have similar brightness, and highly detectable signals have similar brightness. Hence, we may instead expect that species co-occurring should converge for brightness on all patches (prediction 3bis) if the same patches are involved in the same ecological process (communication or camouflage).

Compared to other types of colouration, iridescent colours might enable species recognition on another dimension in the sensory space. Two species can have the same hue or brightness at a given angle but can differ at another angle, via an additional variable we call “hue shift”. Because hue shift cannot be seen at long distances, it may allow species to diverge without interfering with camouflage against predators [24, 90]. Accordingly, we predict overdispersion for hue shift not only on ventral patches, but also on dorsal patches (prediction 4). However, hue shift is often highly correlated with hue due to the optics underlying iridescence (Dakin and Montgomerie [17] for example reported *R*^2^ *≥* 0.95 for the correlation between hue and hue shift). We test this correlation with the data from this article and discuss how it may impact our results.

At the community level, we predict that community colour volume (also known as functional richness `FRic` in functional ecology [87]) and brightness range increase with species richness more than expected in a random species assemblage (null model) because co-occurring species would use different colours (hue or brightness) (prediction 5).

Here we test our five predictions by quantifying both iridescent and non-iridescent colours of 189 hummingbird assemblages in Ecuador that include 112 species and span a large variety of habitats, and by assessing the phenotypic structure (clustering, random distribution, overdispersion of colours) and investigate the underlying processes by taking into account species phylogenetic relatedness within these assemblages. Comparing the uncorrected and the phylogenetically-corrected phenotypic structure of hummingbird communities will allow us to identify which mechanisms (character displacement, species sorting with mutual exclusion of similar species, environmental filtering; as detailed in table 1) underlie the community structure of iridescent colours in hummingbirds.

## Materials and methods

All scripts and data used to produce the results and figures from this article are available at https://doi.org/10.5281/zenodo.3355444.

### Community data

Hummingbirds are particularly suited as a study system to explore the possible effect of reproductive character displacement on iridescent colours because (i) they display a large variety of hues [20] and all species harbour some iridescent patches, many of which have a very strong angular dependence, rapidly shifting from e.g. pink to green or black [22, 26] (but note that many hummingbirds species also have non-iridescent, pigmentary, patches), (ii) they belong to a very speciose family whose phylogeny is well established and readily available [48, 55], (iii) they live only in the Americas, especially in the tropics where numerous species can coexist locally [20] (iv) there is an extensive documentation of hybridisation between co-occurring species (see for example [36, 79] for our region of interest), which creates the perfect opportunity to study reproductive interference and (v) almost all species are available in museum collections and their colour can be objectively measured using spectrometric measurements [23].

Presence/absence data for hummingbird assemblages at 189 sites in Ecuador (see map in fig. S3) were compiled from data in peer-reviewed papers and reports from environmental organisations [34]. These sites cover a large variety of elevation ranges (fig. S3) and habitats [34, 69]. This dataset was previously thoroughly reviewed by comparing the observations with the known elevational and geographical ranges of each species [69] and includes observations of 112 of the 132 hummingbirds species found in Ecuador [73].

### Colour measurements and analyses

For each one of the 112 species, we borrowed one adult male in good condition from either the Museum National d’Histoire Naturelle (MNHN) in Paris or the Musée des Confluences, in Lyon (full list in Online Supplementary Information). Previous studies show that even low sampling per species can accurately capture colour characteristics of the species [18]. Additionally, preliminary analyses on an independent dataset of 834 points across 18 hummingbird species, with up to 5 individuals measured by species, showed that intraspecific coefficient of variation (standard deviation divided by the average) of hue is very low (1.69%) but could be higher for brightness (23.18%) (detailed values for each species in table S3). When comparing intra- to interspecific variation, intraspecific however always remains negligible compared to interspecific variation (intraclass coefficient reported in table S3). We ensured that the specimen colouration was representative of the other specimens available in the collections to the human eye. When multiple subspecies were living in the area where presence was recorded, we randomly picked one of them. Whenever possible, we picked specimens collected in Ecuador (88% of the cases), or when not available in neighbouring countries, such as Colombia or North Peru (11% of the cases), as to minimise the effect of regional variability in colour.

We consistently took spectral reflectance measurements on the eight following patches (described in fig. S1): crown, back, rump, tail, throat, breast, belly, wing. We also made additional measurements on patches that visually differed in colouration from these eight main ones, as in Gomez and Théry [33] and Doutrelant, Paquet, Renoult, Grégoire, Crochet, and Covas [25].

We measured reflectance using a setup similar to Meadows, Morehouse, Rutowski, Douglas, and McGraw [57], relying on the use of two separate optical fibres. Light was conducted from an Oceanoptics DH-2000 lamp emitting over the 300-700 nm range of wavelengths to which birds are sensitive [11] to the sample through an illuminating FC-UV200-2-1.5 x 100 optical fibre (named illumination fibre). Light reflected by the sample was then collected by a second identical optical fibre (named collection fibre) and conducted toward an Oceanoptics USB4000 spectrophotometer (used with the SpectraSuite 2.0.162 software). This setup allows for a precise independent rotation of the illumination and the collection fibres, necessary for the measurements of iridescent colours [65]. For more details about the measurement conditions as recommended in White, Dalrymple, Noble, O’Hanlon, Zurek, and Umbers [89], see the supplementary materials (ESM).

For every patch, we recorded a first reflectance spectrum at the position of the fibres which maximised total reflectance. To measure hue angle dependency (iridescence), we then moved both fibres 10° away from the previous position and recorded a second spectrum, as in Meadows, Roudybush, and McGraw [58]. More recent measurement methods revealed that it would be more accurate to keep the angular span between the illumination and collection fibres constant [39]. We however confirmed that this did not impact our results by running our analyses once with all data and once with only data at a given angular span (which represented 94% of the total data). All measurements were performed in a dark room with temperature control. Recorded spectra were normalised by an Avantes WS-1 white standard and a measurement with the lamp shut down (dark reference) and integration times were determined for each sample as to maximise the intensity of the signal without saturating the spectrometer. Final values were averaged over five consecutive measurements and spectra were smoothed using a loess algorithm and interpolated every 1 nm and negative values were set to zero using the R package `pavo` [52].

We analysed spectra using Endler and Mielke [30] model with relative quantum catches *Q*_*i*_ (without Fechner’s law). All birds are tetrachromats and can see light with wavelengths from 300 to 700 nm, which includes ultra-violet light (UV) [66]. But different bird species vary in their sensitivity [63]: some are UV-sensitive (UVS) while others are violet-sensitive (VS). Literature on colour vision in hummingbirds suggests that both types are found within the family (see Chen and Goldsmith [11] and Herrera, Zagal, Diaz, Fernández, Vielma, Cure, Martinez, Bozinovic, and Palacios [45] for UVS species and Ödeen and Håstad [64] for VS species). Because we did not have enough information to compute ancestral states and vision type for all species in our study and because it was found to have little influence in previous studies [21, 33], we ran our analyses as if all species were VS, using the spectral sensitivities of a typical VS bird, *Puffinus pacificus* [43], whose photoreceptor absorbances match closely those reported for hummingbirds [64]. We used different illuminants defined in Endler [29], depending on the habitat of the species described in Stotz, Fitzpatrick, Parker III, and Moskovits [83] (detailed in SI): “large gaps” illumination was used for species living in the canopy while “forest shade” was used for species living in the understory. Hue was a tridimensional variable defined by the position (*x, y* and *z*) of the reflectance spectrum in the tetrahedron representing bird colour vision space [30] and brightness was defined as in Endler and Mielke [30] (perceived intensity of colour, also sometimes referred to as luminance). We ensured that all indices were repeatable (table S1) by measuring twice the same individual and patch on 20 patches and computing the intra-class coefficient (ICC) with the `rptR` R package [82]. We add another variable to describe iridescence: hue shift, defined as the difference between hue at maximum reflectance and hue at 10° away from maximum reflectance, in a similar fashion to Dakin and Montgomerie [17]. Because it is the difference of two tridimensional variables (hue at the position where reflectance was maximum and hue at 10° away), hue shift is tridimensional as well. Dakin and Montgomerie [17] found a high correlation between hue and hue shift at the intraspecific level in the peacock *Pavo cristatus*, we also report a high correlation at the interspecific level in hummingbirds by performing a linear regression in ℝ^3^ between hue and hue shift (*R*^2^ = 0.51, *F*(3; 1372) = 469.7, *p* < 0.0001). New measurement methods have since been developed and propose a new definition for hue shift which is not correlated to hue but they were not available at the time of this study [39].

We analysed the colour volume for each species by measuring the convex hull volume of all colour patches on the bird, as suggested in Stoddard and Prum [81]. We compared the relationship between the colour volume of a community and the number of species within this community relative to a null model (prediction 5) obtained by creating random assemblages from a species pool containing all species from all communities. In other words, actual assemblages are compared to fictional assemblages with exactly the same number of species but no abiotic or biotic constraints on the species composition.

However, the colour volume does not take into account the patch location on the bird body, raising several concerns. First, two species could use the same colour but at different places on their body. They would then look different to an observer but not identified as such in this analysis. Additionally, we expect different evolutionary signals on different patches, that could even each other out, and blur the outcome at the bird level. For these reasons, we also performed our analyses separately for each one of the following eight patches: crown, back, rump, tail, throat, breast, belly, wing (locations shown in fig. S1).

### Trochilidae phylogeny and comparative analyses

A distribution of 100 phylogenetic trees of the Trochilidae family was downloaded from birdtree.org [48] to take into account phylogenetic uncertainty in the comparative analyses [67]. The 112 species included in this study constitute a fairly even sampling of the hummingbird phylogeny (fig. S2).

We used the method developed by Hardy and Senterre [42] and Baraloto, Hardy, Paine, Dexter, Cruaud, Dunning, Gonzalez, Molino, Sabatier, Savolainen, and Chave [5] to analyse respectively the phylogenetic (Π_*ST*_) and phenotypic (*τ*_*ST*_) structures of the hummingbird communities of Ecuador (clustering or overdispersion). This method relies on computing indices inspired by the Simpson index and the fixation index *F*_*ST*_, comparing the observed diversity within and between communities. For phylogeny, Π_*ST*_ can reveal phylogenetic clustering (Π_*ST*_ > 0) or phylogenetic overdispersion (Π_*ST*_ < 0) within communities. Likewise, for phenotypic traits, *τ*_*ST*_ can reveal phenotypic clustering (*τ*_*ST*_ > 0) or phenotypic overdispersion (*τ*_*ST*_ < 0) within communities. Statistical significance of overdispersion or clustering is obtained from comparing the observed value to that obtained for the same patch location from 1000 random communities (created by drawing from the total species pool, using algorithm 1s from Hardy [41], which keeps the local species richness per site constant). This approach compares the phenotypic structure to what would be expected by chance.

To disentangle the relative effect of ecological (species sorting) and evolutionary mechanisms (selection), we also perform our analyses by taking into account the phylogenetic relationships between species. If the species in the community are more clustered or overdispersed than expected given their phylogenetic relationships, this is taken as evidence that the trait has not evolved in a Brownian fashion (detailed in table 1). To this end, we used the `decouple` function [19], which returns phylogenetically predicted and residual trait values by performing a linear regression of individual trait values explained by the phylogeny. We computed the value of *τ*_*ST*_ on trait values decoupled from the phylogeny. This value is hereafter denoted *dcτ*_*ST*_. Similarly to the classical *τ*_*ST*_, the sign of *dcτ*_*ST*_ indicates phenotypic clustering (*dcτ*_*ST*_ > 0) or overdispersion (*dcτ*_*ST*_ < 0) once the effect of the phylogenetic structure of the communities has been decoupled.

Analyses performed on a tree distribution (Π_*ST*_ and *dcτ*_*ST*_) with *n* trees return a distribution of *n* statistics values and *n* p-values *p*_*i*_. We summarised this information by computing the median of the statistics and the overall p-value *p* by using Jost’s formula [4]: 

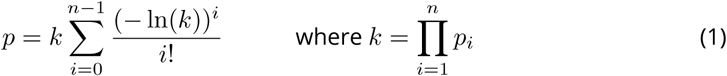

## Results

We find a strong phylogenetic clustering within communities (Π_*ST*_ = 0.062 > 0, *p* < 0.0001), indicating that co-occurring species are more closely related than expected by chance.

### Phenotypic structure of the communities (predictions 1 - 4)

When looking at the bird entire body (when all patches are included simultaneously) by computing the overlap of the colour volumes, we did not find any phenotypic structure.

When the different major patches (crown, back, rump, tail, throat, breast, belly and wing) are examined separately (table 2 and table S2), we find clustering (*τ*_*ST*_ > 0) in hue and hue shift on the back, rump, tail, belly and wing. Once we decouple the effect of the shared evolutionary history, we find clustering on the crown and the back (*dcτ*_*ST*_ > 0) but overdispersion on the belly for both hue and hue shift (*dcτ*_*ST*_ < 0). Hue shift is also overdispersed on the rump and the tail (*dcτ*_*ST*_ < 0). There is no phenotypic structure on the throat, breast or wing for hue and hue shift nor on the rump or the tail for hue.

**Table 2.**
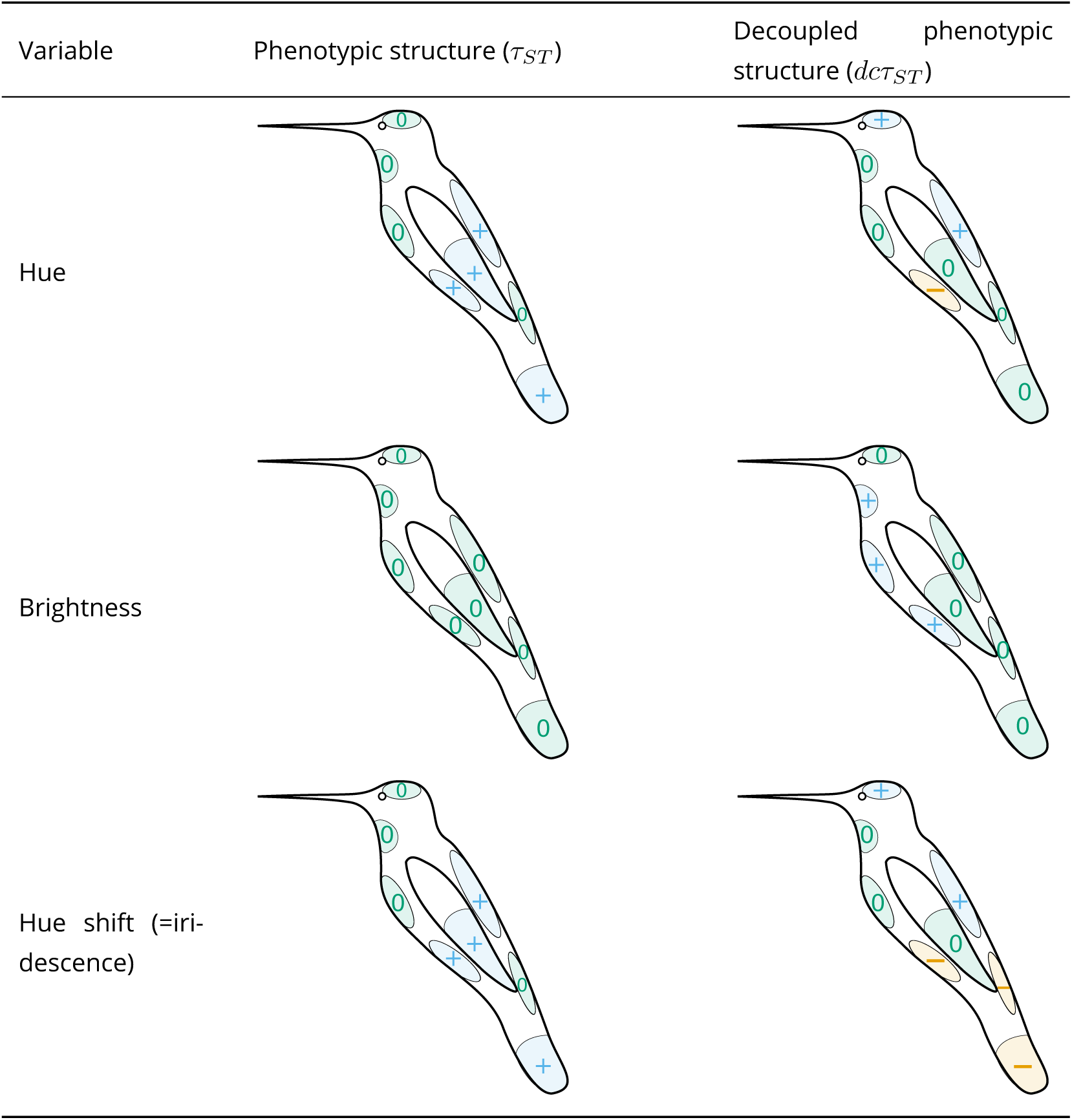
Phenotypic structure of hummingbird communities for different variables (hue, brightness and hue shift) on the patches studied (crown, back, rump, tail, throat, breast, belly, wing; names and locations illustrated in fig. S1). Hue is a tridimensional variable defined by the reflectance spectrum position *x, y* and *z* in the tetrahedron representing avian colour space. Blue plus signs + indicate significant phenotypic clustering (*τ*_*ST*_ or *dcτ*_*ST*_ > 0), orange minus signs − indicate significant phenotypic overdispersion (*τ*_*ST*_ or *dcτ*_*ST*_ < 0), and green zeros 0 represent the absence of phenotypic structure. The left column shows the raw phenotypic structure of the community (columns in table 1), which may be influenced by the phylogenetic structure while the right column shows the phenotypic structure of the community, decoupled from all effects caused by the phylogeny (rows in table 1). By comparing the values of *τ*_*ST*_ and *dcτ*_*ST*_ for each trait colour variable (hue, brightness and hue shift), we can assume a probable evolutionary scenario for each patch, based on the explanation in table 1. Exact values for the statistics are available in table S2.

We find no phenotypic structure (neither clustering nor overdispersion) for brightness on any patches before phylogenetic correction. After phylogenetic correction, brightness values for the throat, breast and belly are clustered among co-occurring species (*dcτ*_*ST*_ > 0) but show no phenotypic structure for the crown, the back, the wing and the tail.

### Effect of community species richness on colour characteristics (prediction 5)

We found that the brightness range within a community increased in the same way as a null model built from random species assemblages (fig. 1b). For colour volume, we find some outliers with a higher colour volume than expected for community with the same number of species (fig. 1a).

**Figure 1.**
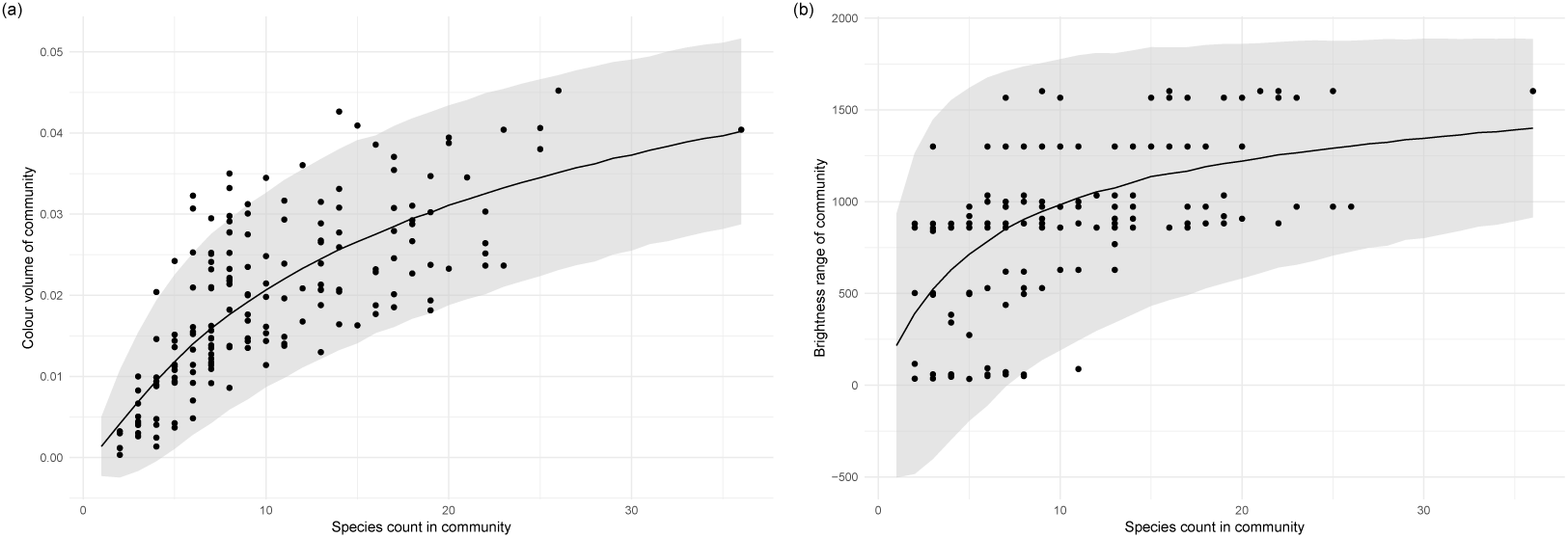
(a) Community total colour volume and (b) brightness range increase with the number of species within the community. Each point is a community. The black solid line represents the mean value of (a) colour volume or (b) brightness range from 10 000 random communities with a given species count (null model) and the gray ribbon represents two standard deviations from the mean of the null model.

## Discussion

Our findings are consistent with our hypothesis that colour structure within hummingbird communities likely results from the interplay between two selective pressures, acting in opposite directions: selection by the local environment (e.g. camouflage from predators, leading to phenotypic clustering on dorsal patches, and selection for species recognition, leading to phenotypic overdispersion on ventral and facial patches. We also discuss other possible effects that might have contributed to the observed pattern.

### Evidence for different evolutionary scenarios depending on patch location

At the entire bird level (i.e. when pooling together all patches), we did not find any phenotypic structure. But as mentioned earlier, this was expected since different locations on the birds are thought to be under different selection regimes [21, 33].

In accordance with our prediction 5, community colour volume (as estimated by the convex hull of hue and brightness range within a community) increases slightly faster with the number of species in the community than predicted by a null model. This suggests that co-occurring species in these communities tend to use more similar colours than expected by chance. However, this is not the case for the majority of communities, where co-occurring species do not use more nor less similar colours than expected by chance. This is further confirmed by the absence of phenotypic structure on the colour volume and the brightness when the effect of the phylogeny is not decoupled.

This could be the consequence of similar selective pressures between the communities we studied, leading colours in all assemblages to be randomly determined. This is however not very likely because the communities we studied differ a lot in both their vegetation background and therefore in the pressure for crypsis [34] and in their species composition. A more likely hypothesis is that co-occurring species tend to use the same colours but not necessarily on the same patches, which would also explain the absence of phenotypic structure when we pool all patches without taking into account their location. This is confirmed by our analysis patch by patch, where we find either clustering or overdispersion depending on the location of the patch.

### Selection for convergence and phenotypic clustering

In accordance with our first two predictions, co-occurring hummingbird species tend to have similar hues on patches more likely dedicated to camouflage (back, rump, tail, wing; prediction 1) but not on patches more likely used in communication (crown, throat, breast; prediction 2), as shown in table 2 and table S2. This new result for iridescent colours matches what has been previously described for non-iridescent colours [21, 33]. The phenotypic clustering observed for hue on the rump, the tail and the wing vanishes after decoupling the clustering effect due to phylogenetic structure. This suggests that phenotypic clustering of hue on the rump, the tail and the wing is not caused by convergent evolution of co-occurring species but by environmental filtering, leading related, similar-looking species to live in the same area (as explained in table 1). This is confirmed by the high value of phylogenetic clustering. This sign of phylogenetic clustering complements the results from Graham, Parra, Rahbek, and McGuire [34] on the same dataset. We showed that intra-community species relatedness is high compared to inter-community species relatedness (Π_*ST*_), while they showed that intra-community species relatedness (Net Relatedness Index) is higher than expected from random assemblages in 71% of the cases [34]. This phylogenetic clustering may be caused by a strong niche conservatism but our study cannot discriminate whether such niche conservatism involves colour or other ecological traits. Our data does not allow us to assert with certainty the evolutionary history from the pattern we observe but the predominance of green and brown hues on the back and the wing respectively, as shown in fig. S4, hints to a role in camouflage. Alternatively, this phylogenetic clustering could be caused by hummingbirds’ costly hovering flight at high elevation due to weaker lift caused by the decreasing atmospheric pressure [1, 2, 84], high foraging specialisation [49] or low dispersal ability, but this last hypothesis remains quite unlikely as the rare studies on this topic have shown that different hummingbird species display a wide variation in their dispersal ability [10, 60].

Contrary to our prediction 2, we also find clustering of hue on the belly before the use of the `decouple` function. However, the fact that it turns into overdispersion after the use of the `decouple` function, and not simply into a random phenotypic structure (as opposed to the rump, the tail and the wing mentioned just before), suggests this initial clustering (right column in table 1) is mainly caused by environmental filtering on another trait but that hue on the belly is still under selection for divergence (first row in table 1). This other trait may be the colour of another patch or other ecological traits, as we explained previously.

We found a significant clustering of brightness on the throat, breast and belly after controlling for the phylogeny, indicating that brightness on those patches is more similar than expected given the phylogeny among co-occurring species (prediction 3bis). This suggests that the same patches have been selected to be involved either in communication or in camouflage among species living in the same environment. This is seen after controlling for the phylogeny and it is therefore not caused by the phylogenetic relatedness of co-occurring species. This is not surprising as many studies showed the paramount importance of the throat in the courtship display of many hummingbird species [46, 75–78] Two main hypotheses can explain why co-occurring species tend to communicate (or camouflage themselves) using the same patches: (i) There may be selective pressures for the use of specific patches in camouflage in a given environment (e. g., patches that are more exposed to predators’ sight). (ii) Convergence in patches used in communication may be selected because it improves competitor identification in the case of a strong ecological niche overlap (convergence by agonistic character displacement as shown in Grether, Losin, Anderson, and Okamoto [37] and Tobias, Planqué, Cram, and Seddon [85]).

All those results suggest a strong effect of the environment in the evolution of colour in agreement with McNaught and Owens [56] who found that bird plumage colour was due to the light environment and not to reproductive character displacement in Australian birds. However, we do not find clustering on all patches, which suggests that, for some patches, the effect of habitat pressure is somehow limited or counterbalanced by reproductive or agonistic character displacement. On the contrary, for some patches, we found patterns that are likely the result of character displacement.

### Character displacement and phenotypic overdispersion

In agreement with our prediction 2, after decoupling the effect of the phylogeny, there is overdispersion of hue on the belly, likely caused by character displacement (table 1). At a completely different taxonomic scale, focusing on a single hummingbird genus (*Coeligena*) with 11 species, Parra [70] also found that the belly was always involved in the difference in hue between subspecies. It was sometimes even the only patch causing those differences, as for example between *Coeligena torquata fulgidigula* and *Coeligena torquata torquata*. This suggests that the interspecific divergence we found on the belly at the community level on the whole Trochilidae family can be observed at different geographic and taxonomic scales, and even between subspecies of the same species.

As predicted, we also find more phenotypic overdispersion for hue shift than hue after decoupling the effect of the phylogeny, for example, on the rump and on the tail (prediction 4). It is possible that hue shift is less sensitive to selection for convergence because it may vary without disturbing camouflage efficacy. However, we did not find the expected relaxing of clustering on hue shift on patches such as the back. This is likely caused by the fact that hue shift is highly correlated with hue, as found in this study and in Dakin and Montgomerie [17], who used the same indices to quantify iridescence. This correlation is due to the optics controlling iridescence, meaning that species that display similar hues should also display the same hue shift if they use the same underlying multilayer structures. The fact that the correlation is not perfect and that we nonetheless get different phenotypic patterns for hue and hue shift on some patches suggests that co-occurring species use different multilayer structures (as recently confirmed by [40]), which can produce different iridescent effects while displaying the same hue (functional convergence on hue).

Against our prediction 2, we did not find phenotypic overdispersion on any of the colour variables on patches such as the throat or the crown, that are thought to be sexually selected and often used in courtship displays [15, 78]. Several hypotheses can explain this fact: (i) The overdispersion on some patches (hue on the belly and hue shift on the rump and tail) is sufficient to enable species recognition. (ii) The current phenotypic structure, which is neither overdispersed nor clustered, on those patches is sufficient to enable species recognition. Indeed, the absence of phenotypic overdispersion does not mean that species look the same. It simply means that colour differences between species living in the same community and species in different communities occur in similar ranges. This difference may be sufficient to relax the selective pressure towards reproductive character displacement. (iii) The pressure towards overdispersion is balanced by habitat filtering (for both ventral and dorsal patches), resulting in no apparent phenotypic structure. The latter hypothesis was also a candidate explanation of the pattern found by Martin, Montgomerie, and Lougheed [53], where sympatric closely related species are more divergent than allopatric ones, but only when the range overlap is limited. They suggested that local adaptation could hinder divergence when species ranges was exactly the same.(iv) Species recognition is achieved by additional means and divergence occurs on others traits, such as modified feathers [28], song [51, 54] or non-vocal noises [12–14] and size. Notably, different species of hummingbirds can have very different courtship behaviour: leks for hermits [71, 80], dives and shuttle displays for bees [13, 47, 77], for instance.

Taken together, our results suggest that hummingbird iridescent colours are determined by different evolutionary mechanisms depending on their location. Within a community, co-occurring hummingbird species tend to display the same hues on dorsal patches which is what we expect if colour on these patches is mainly driven by selective pressures related to the local environment, such as selection for crypsis by predators, causing phenotypic clustering at the community level. This phenotypic clustering does not seem to be caused by adaptive convergence on colours but rather by environmental filtering perhaps linked to other ecological traits such as elevation tolerance or flight ability. In spite this suspected environmental filtering, there is overdispersion for hue on the belly and hue shift on the rump and the tail. This suggest a possible role of character displacement, which could mean that iridescence could be used a way to enable species recognition without affecting camouflage efficacy of birds, by opening up a new dimension in the sensory space: hue shift.

## Data accessibility

Data are available online: https://doi.org/10.5281/zenodo.3355443

## Acknowledgments

This project heavily relied on museum specimens which were made available by the work of collection curators: Patrick Boussès, Anne Previato, and Jérôme Fuchs (Muséum National d’Histoire Naturelle), Cédric Audibert and Harold Labrique (Musée des Confluences). J.L.P was funded by a Colombian Administrative Department for Science and Technology – Colciencias Grant code 111571250482 – contract number 248-2016. We also thank PCI recommender Sebastien Lavergne and two anonymous reviewers for their comments which helped us improve this manuscript. Version 5 of this preprint has been peer-reviewed and recommended by Peer Community In Evolutionary Biology (https://doi.org/10.24072/pci.evolbiol.100086).

## Conflict of interest disclosure

The authors of this preprint declare that they have no financial conflict of interest with the content of this article. Marianne Elias is part of the managing board of PCIEvolBiol and is one of the PCIEvolBiol recommenders.

## Appendix

**Table 3.**
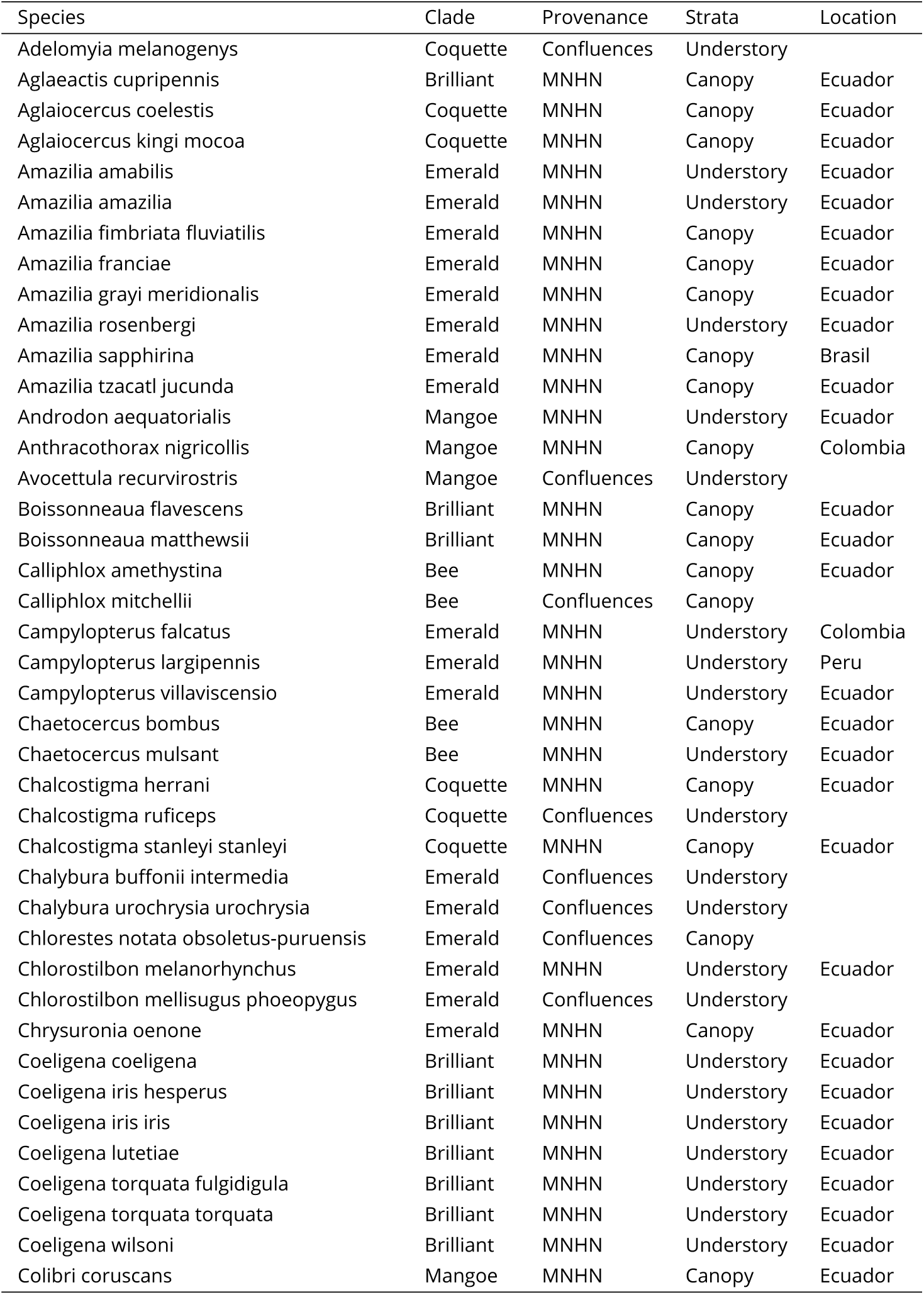
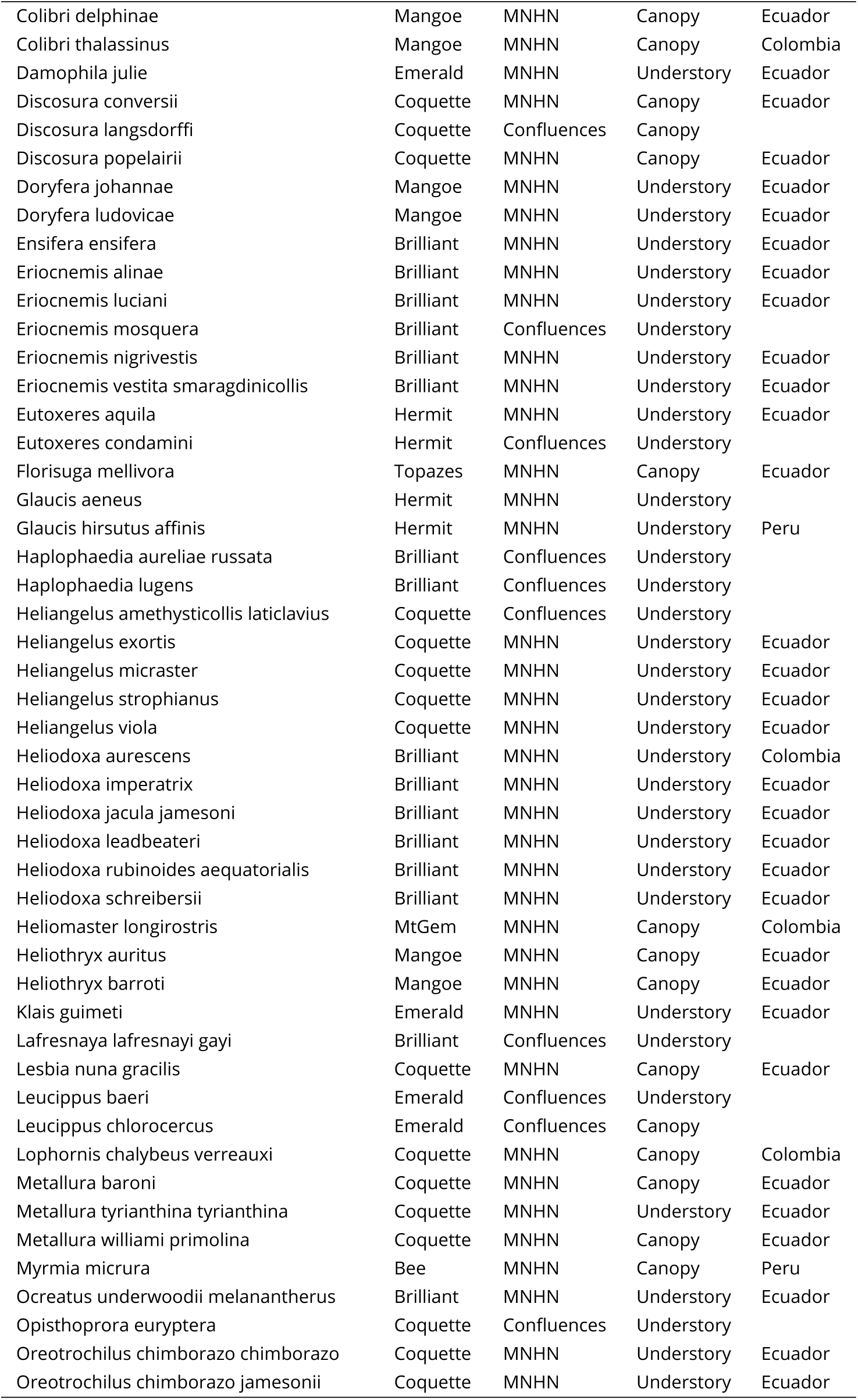
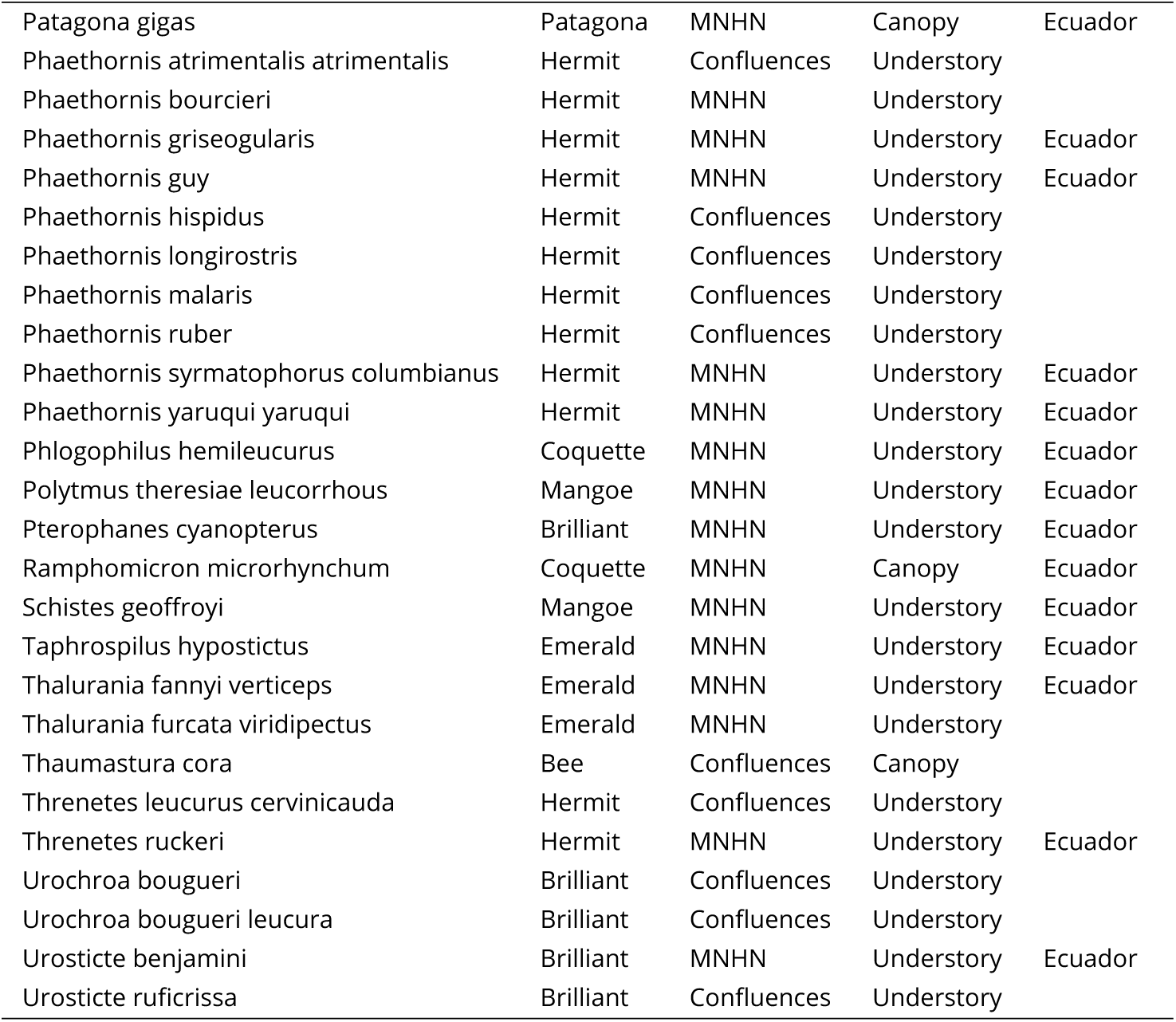
List of species with their provenance (Confluences = Musée des Confluences, Lyon, France, MNHN = Muséum National d’Histoire Naturelle, Paris, France), strata, and place of collection (when known). Strata data were extracted from Stotz, Fitzpatrick, Parker III, and Moskovits [83] and used in vision models.

| Species | Clade | Provenance | Strata | Location |
| --- | --- | --- | --- | --- |
| *Adelomyia melanogenys* | Coquette | Confluences | Understory |  |
| *Aglaeactis cupripennis* | Brilliant | MNHN | Canopy | Ecuador |
| *Aglaiocercus coelestis* | Coquette | MNHN | Canopy | Ecuador |
| *Aglaiocercus kingi mocoa* | Coquette | MNHN | Canopy | Ecuador |
| *Amazilia amabilis* | Emerald | MNHN | Understory | Ecuador |
| *Amazilia amazilia* | Emerald | MNHN | Understory | Ecuador |
| *Amazilia fimbriata fluviatilis* | Emerald | MNHN | Canopy | Ecuador |
| *Amazilia franciae* | Emerald | MNHN | Canopy | Ecuador |
| *Amazilia grayi meridionalis* | Emerald | MNHN | Canopy | Ecuador |
| *Amazilia rosenbergi* | Emerald | MNHN | Understory | Ecuador |
| *Amazilia sapphirina* | Emerald | MNHN | Canopy | Brasil |
| *Amazilia tzacatl jucunda* | Emerald | MNHN | Canopy | Ecuador |
| *Androdon aequatorialis* | Mangoe | MNHN | Understory | Ecuador |
| *Anthracothonax nigricollis* | Mangoe | MNHN | Canopy | Colombia |
| *Avocettula recurvirostris* | Mangoe | Confluences | Understory |  |
| *Boissonneaua flavescens* | Brilliant | MNHN | Canopy | Ecuador |
| *Boissonneaua matthewsii* | Brilliant | MNHN | Canopy | Ecuador |
| *Calliphlox amethystina* | Bee | MNHN | Canopy | Ecuador |
| *Calliphlox mitchellii* | Bee | Confluences | Canopy |  |
| *Campylopterus falcatus* | Emerald | MNHN | Understory | Colombia |
| *Campylopterus largipennis* | Emerald | MNHN | Understory | Peru |
| *Campylopterus villaviscensio* | Emerald | MNHN | Understory | Ecuador |
| *Chaetocercus bombus* | Bee | MNHN | Canopy | Ecuador |
| *Chaetocercus mulsant* | Bee | MNHN | Understory | Ecuador |
| *Chalcostigma herrani* | Coquette | MNHN | Canopy | Ecuador |
| *Chalcostigma ruficeps* | Coquette | Confluences | Understory |  |
| *Chalcostigma stanleyi stanleyi* | Coquette | MNHN | Canopy | Ecuador |
| *Chalybura buffonii intermedia* | Emerald | Confluences | Understory |  |
| *Chalybura urochrysis urochrysis* | Emerald | Confluences | Understory |  |
| *Chlorestes notata obsoletus-puruensis* | Emerald | Confluences | Canopy |  |
| *Chlorostilbon melanorhynchus* | Emerald | MNHN | Understory | Ecuador |
| *Chlorostilbon mellisugus phoeopygus* | Emerald | Confluences | Understory |  |
| *Chrysuronia oenone* | Emerald | MNHN | Canopy | Ecuador |
| *Coeligena coeligena* | Brilliant | MNHN | Understory | Ecuador |
| *Coeligena iris hesperus* | Brilliant | MNHN | Understory | Ecuador |
| *Coeligena iris iris* | Brilliant | MNHN | Understory | Ecuador |
| *Coeligena lutetiae* | Brilliant | MNHN | Understory | Ecuador |
| *Coeligena torquata fulgidigula* | Brilliant | MNHN | Understory | Ecuador |
| *Coeligena torquata torquata* | Brilliant | MNHN | Understory | Ecuador |
| *Coeligena wilsoni* | Brilliant | MNHN | Understory | Ecuador |
| *Colibri coruscans* | Mangoe | MNHN | Canopy | Ecuador |
| Colibri delphinae | Mangoe | MNHN | Canopy | Ecuador |
| Colibri thalassinus | Mangoe | MNHN | Canopy | Colombia |
| Damophila julie | Emerald | MNHN | Understory | Ecuador |
| Discosura conversii | Coquette | MNHN | Canopy | Ecuador |
| Discosura langsdorffi | Coquette | Confluences | Canopy |  |
| Discosura popelairii | Coquette | MNHN | Canopy | Ecuador |
| Doryfera johannae | Mangoe | MNHN | Understory | Ecuador |
| Doryfera ludovicae | Mangoe | MNHN | Understory | Ecuador |
| Ensifera ensifera | Brilliant | MNHN | Understory | Ecuador |
| Eriocnemis alinae | Brilliant | MNHN | Understory | Ecuador |
| Eriocnemis luciani | Brilliant | MNHN | Understory | Ecuador |
| Eriocnemis mosquera | Brilliant | Confluences | Understory |  |
| Eriocnemis nigrivestis | Brilliant | MNHN | Understory | Ecuador |
| Eriocnemis vestita smaragdinicollis | Brilliant | MNHN | Understory | Ecuador |
| Eutoxeres aquila | Hermit | MNHN | Understory | Ecuador |
| Eutoxeres condomini | Hermit | Confluences | Understory |  |
| Florisuga mellivora | Topazes | MNHN | Canopy | Ecuador |
| Glaucis aeneus | Hermit | MNHN | Understory |  |
| Glaucis hirsutus affinis | Hermit | MNHN | Understory | Peru |
| Haplophaedia aureliae russata | Brilliant | Confluences | Understory |  |
| Haplophaedia lugens | Brilliant | Confluences | Understory |  |
| Heliangelus amethysticollis laticlavus | Coquette | Confluences | Understory |  |
| Heliangelus exortis | Coquette | MNHN | Understory | Ecuador |
| Heliangelus micraster | Coquette | MNHN | Understory | Ecuador |
| Heliangelus strophianus | Coquette | MNHN | Understory | Ecuador |
| Heliangelus viola | Coquette | MNHN | Understory | Ecuador |
| Heliodoxa aurescens | Brilliant | MNHN | Understory | Colombia |
| Heliodoxa imperatrix | Brilliant | MNHN | Understory | Ecuador |
| Heliodoxa jacula jamesoni | Brilliant | MNHN | Understory | Ecuador |
| Heliodoxa leadbeateri | Brilliant | MNHN | Understory | Ecuador |
| Heliodoxa rubinoides aequatorialis | Brilliant | MNHN | Understory | Ecuador |
| Heliodoxa schreibersii | Brilliant | MNHN | Understory | Ecuador |
| Helioaster longirostris | MtGem | MNHN | Canopy | Colombia |
| Heliothryx auritus | Mangoe | MNHN | Canopy | Ecuador |
| Heliothryx barroti | Mangoe | MNHN | Canopy | Ecuador |
| Klais guimeti | Emerald | MNHN | Understory | Ecuador |
| Lafresnaya lafresnaya gayi | Brilliant | Confluences | Understory |  |
| Lesbia nuna gracilis | Coquette | MNHN | Canopy | Ecuador |
| Leucippus baeri | Emerald | Confluences | Understory |  |
| Leucippus chlorocercus | Emerald | Confluences | Canopy |  |
| Lophornis chalybeus verreauxi | Coquette | MNHN | Canopy | Colombia |
| Metallura baroni | Coquette | MNHN | Canopy | Ecuador |
| Metallura tyrianthina tyrianthina | Coquette | MNHN | Understory | Ecuador |
| Metallura williamsi primolina | Coquette | MNHN | Canopy | Ecuador |
| Myrmia micrura | Bee | MNHN | Canopy | Peru |
| Ocreatus underwoodii melanatherus | Brilliant | MNHN | Understory | Ecuador |
| Opisthoprora euryptera | Coquette | Confluences | Understory |  |
| Oreotrochilus chimborazo chimborazo | Coquette | MNHN | Understory | Ecuador |
| Oreotrochilus chimborazo jamesonii | Coquette | MNHN | Understory | Ecuador |
| Patagona gigas | Patagona | MNHN | Canopy | Ecuador |
| Phaethornis atrimentalis atrimentalis | Hermit | Confluences | Understory |  |
| Phaethornis bourcieri | Hermit | MNHN | Understory |  |
| Phaethornis griseogularis | Hermit | MNHN | Understory | Ecuador |
| Phaethornis guy | Hermit | MNHN | Understory | Ecuador |
| Phaethornis hispidus | Hermit | Confluences | Understory |  |
| Phaethornis longirostris | Hermit | Confluences | Understory |  |
| Phaethornis malaris | Hermit | Confluences | Understory |  |
| Phaethornis ruber | Hermit | Confluences | Understory |  |
| Phaethornis syrmatophorus columbianus | Hermit | MNHN | Understory | Ecuador |
| Phaethornis yaruqui yaruqui | Hermit | MNHN | Understory | Ecuador |
| Phlogophilus hemileucurus | Coquette | MNHN | Understory | Ecuador |
| Polytmus theresiae leucorrhous | Mangoe | MNHN | Understory | Ecuador |
| Pterophanes cyanopterus | Brilliant | MNHN | Understory | Ecuador |
| Ramphomicron microrhynchum | Coquette | MNHN | Canopy | Ecuador |
| Schistes geoffroyi | Mangoe | MNHN | Understory | Ecuador |
| Taphrospilus hypostictus | Emerald | MNHN | Understory | Ecuador |
| Thalurania fannyi verticeps | Emerald | MNHN | Understory | Ecuador |
| Thalurania furcata viridipectus | Emerald | MNHN | Understory |  |
| Thaumastura cora | Bee | Confluences | Canopy |  |
| Threnetes leucurus cervinicauda | Hermit | Confluences | Understory |  |
| Threnetes ruckeri | Hermit | MNHN | Understory | Ecuador |
| Urochroa bougueri | Brilliant | Confluences | Understory |  |
| Urochroa bougueri leucura | Brilliant | Confluences | Understory |  |
| Urosticte benjamini | Brilliant | MNHN | Understory | Ecuador |
| Urosticte ruficrissa | Brilliant | Confluences | Understory |  |

**Supplementary figure 1.**
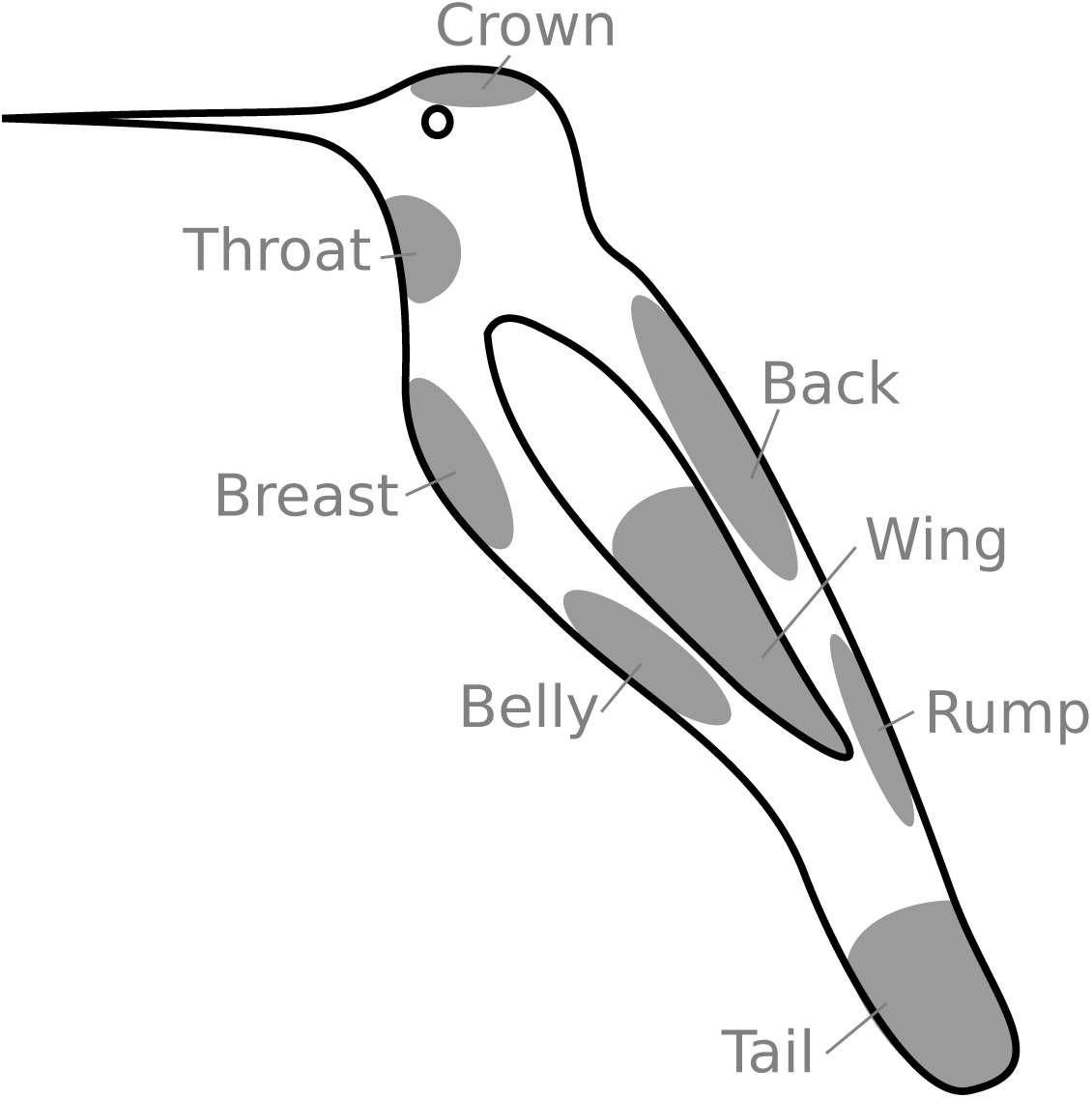
Locations and names of the 8 patches measured on all species. Additional patches were measured for each species as soon as they differed from one of the 8 patches listed here for a human observer, as detailed in the methods section and as in Gomez and Théry [33].

**Supplementary figure 2.**
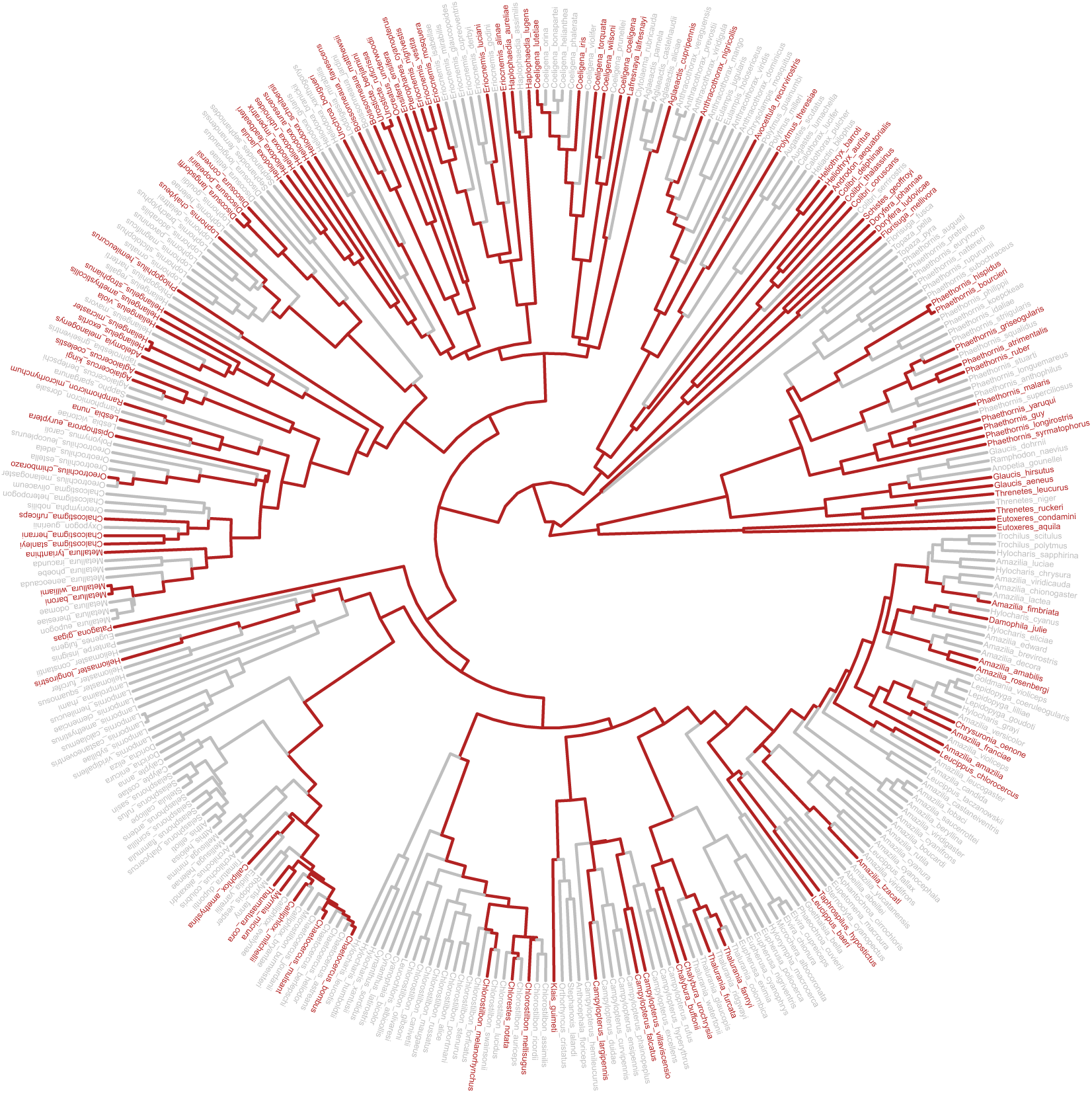
Phylogenetic coverage of the *Trochilidae* family in our dataset (species and lineages in red).

**Supplementary figure 3.**
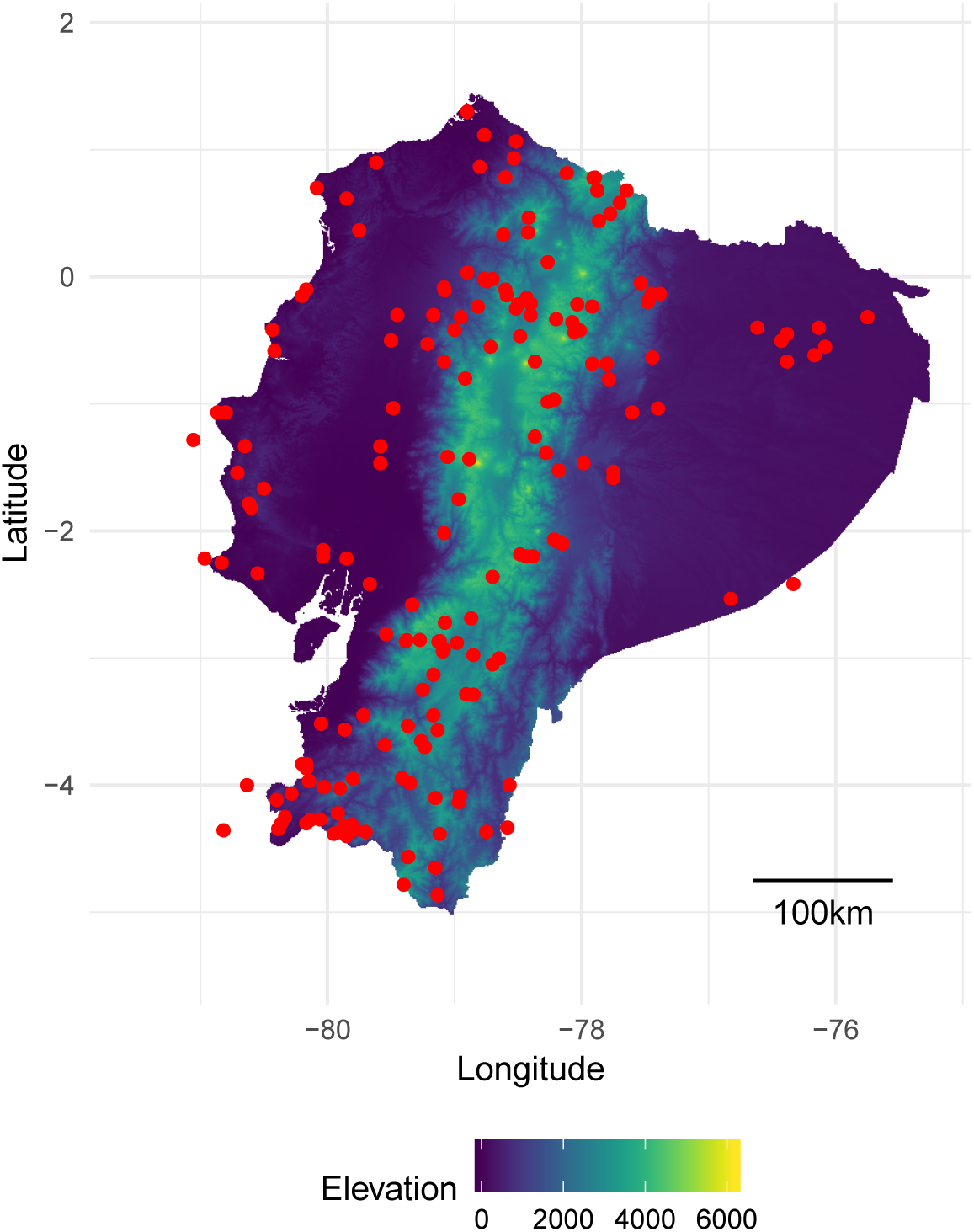
Study site locations (red dots) plotted on an altitudinal map of Ecuador. Communities outside the borders of the map are on islands or close enough to Ecuador borders to be taken into account in our study.

**Table 4.** Measurement of intraspecific variability for brightness (B2) and hue (H1) by computing the coefficient of variation (standard deviation divided the average) on an independent dataset of hummingbirds living in French Guiana (Gomez *et al*, unpublished data), in which between 2 and 5 males (last column) were measured for each species. The measurement protocol differs slightly from the one used in this study, because we used a birfucated probe at 45°, which may increase the intraspecific variability in brightness. In spite of the apparently high values of the coefficient of variation for brightness, it remains highly repeatable as estimated by the intra-class coefficient [62]: *R* = 0.809, *p* < 0.0001 for brightness and *R* = 0.661, *p* < 0.0001 for hue.

| Species | CV brightness (%) | CV_ hue (%) | n |
| --- | --- | --- | --- |
| Anthracothonax nigricollis | 20.57 | 2 | 3 |
| Calliphlox amethystina | 24.37 | 1.13 | 5 |
| Campylopterus largipennis | 17.43 | 0.1 | 2 |
| Chlorestes notatus | 19.79 | 1.96 | 5 |
| Discosura longicauda | 26.27 | 2.51 | 5 |
| Florisuga mellivora | 22.41 | 2.1 | 5 |
| Glaucis hirsuta | 33.75 | 0 | 4 |
| Helimaster longirostris | 26.88 | 2.26 | 4 |
| Heliothryx aurita | 22.82 | 1.26 | 5 |
| Hylocharis cyanus | 29.75 | 2.55 | 3 |
| Hylocharis sapphirina | 23.32 | 3.36 | 4 |
| Lophornis ornatus | 23.38 | 1.55 | 5 |
| Phaethornis longuemareus | 18.59 | 0.15 | 4 |
| Phaethornis malaris | 21.44 | 0.1 | 2 |
| Phaethornis superciliosus | 27.88 | 0.1 | 5 |
| Thalurania furcata | 84.13 | 12.4 | 2 |
| Threnetes niger | 16.42 | 0.1 | 2 |
| Topaza pella | 23.04 | 1.83 | 5 |

**Supplementary table 1.**
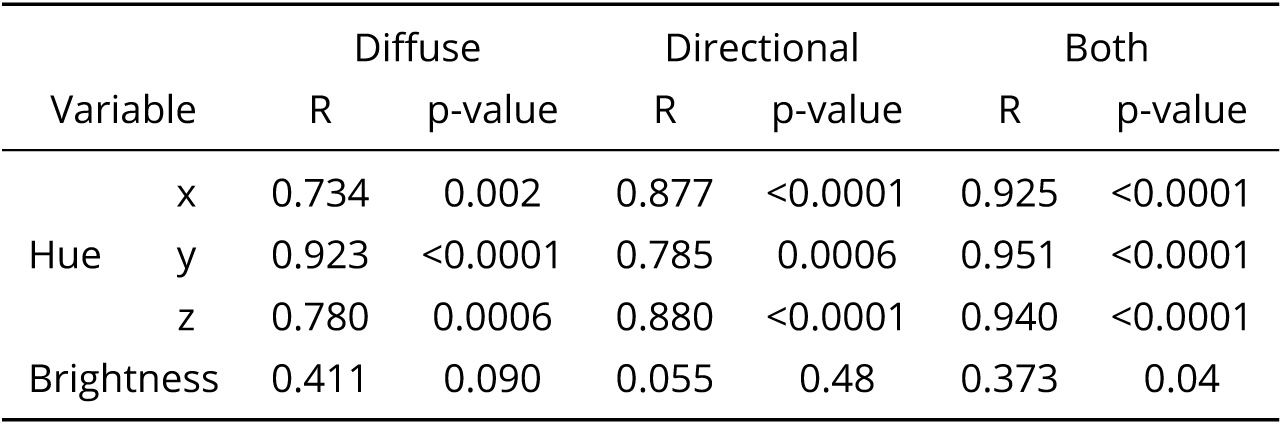
We quantified the repeatability R (intra-class coefficient ICC) and the related p-value by boostraping using the `rptR` R package [62] of indices used in this study by performing the same measurements twice on two patches for 12 species (*Coeligena torquata, Colibri coruscans, Doryfera ludovicae, Heliangelus strophianus, Heliodoxa jamesonii, Heliothryx barroti, Juliamyia julie, Lesbia nuna, Metallura tyrianthina, Ramphomicron microrhynchum, Schistes albogularis, Urosticte benjamini*). Patches were selected to be of similar hue from a human point of view.

**Supplementary figure 4.**
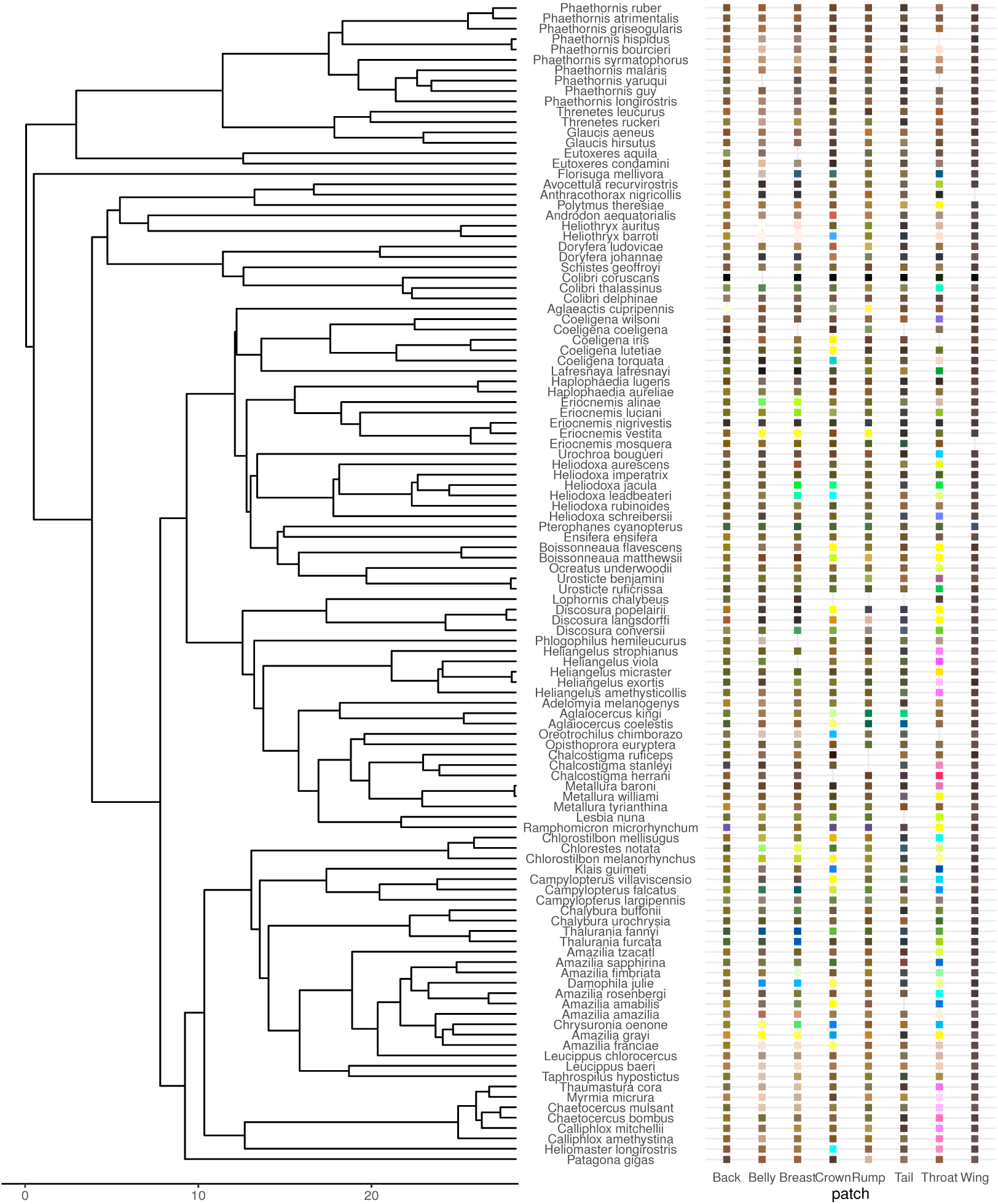
Colour of the 8 main patches for each species in our dataset. The colour corresponds to the colour in the human visual system (CIE10). The x-axis on the phylogeny is in millions years.

**Supplementary table 2.**
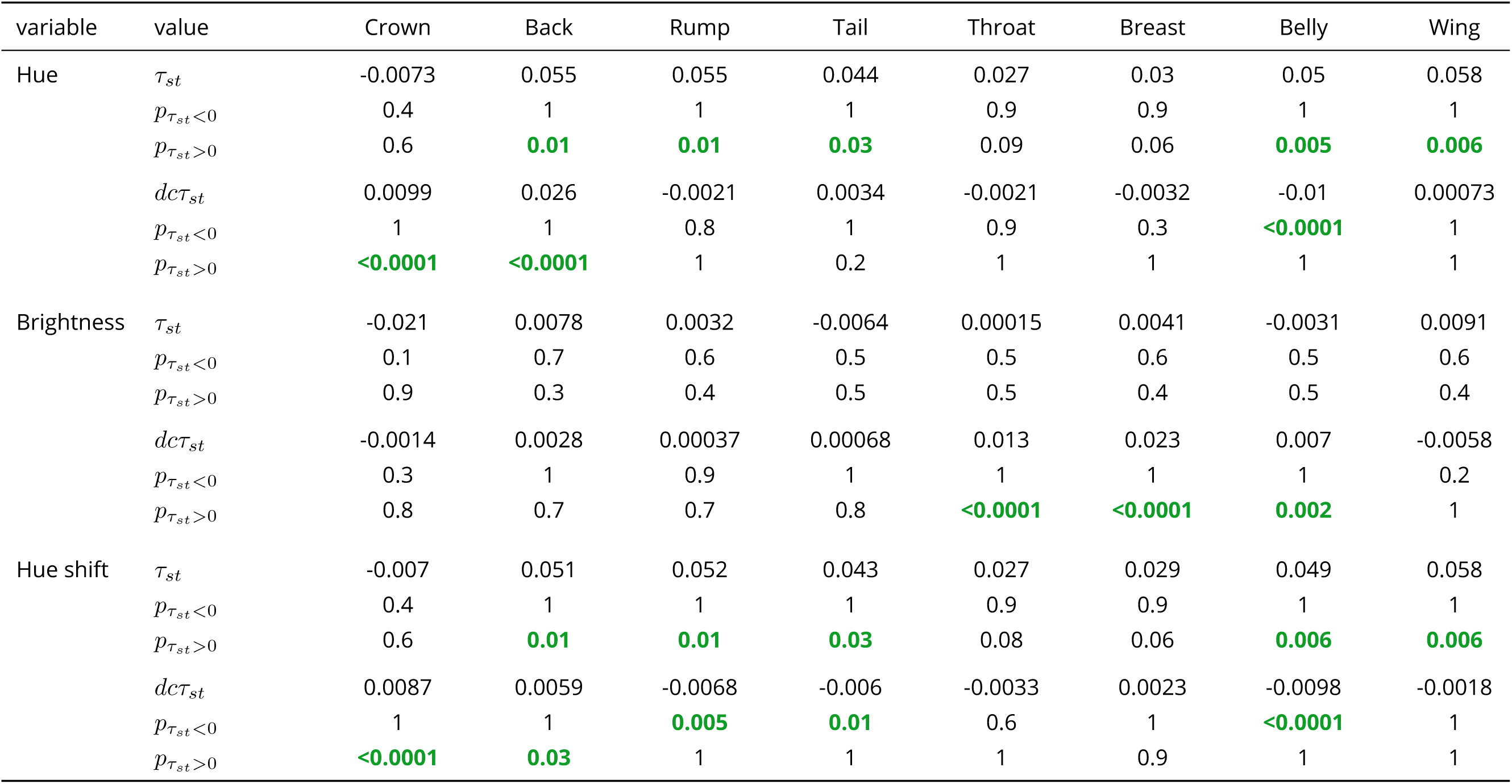
Numerical values for *τ*_*st*_ and decoupled *τ*_*st*_ (denoted *dcτ*_*st*_). P-values were computed by comparison of the actual value with the null distribution (obtained by randomisation of the communities using method `1s` of Hardy [41]). Significant p-values are in bold and green. Positive values of *dcτ*_*st*_ indicate phenotypic clustering whereas negative values indicate overdispersion.

## Notes

#### Summary of Updates

Final (recommended) version for PCI EvolBiol

https://doi.org/10.5281/zenodo.3355443

